# Ryanodine receptor cluster remodeling links *COL4A2* mutation to cerebral artery dysfunction

**DOI:** 10.64898/2026.09.25.754477

**Authors:** Boris Lavanderos, Evan Yamasaki, Alfredo Sanchez-Solano, Wanchun Zhu, Cassandre Labelle-Dumais, Yumei Feng Earley, Douglas B. Gould, Scott Earley

## Abstract

Pathogenic variants of *COL4A1* and *COL4A2*, which encode the α1 and α2 chains of type IV collagen in vascular basement membranes, cause cerebral small vessel disease and intracerebral hemorrhage, but the signaling mechanisms that precede vascular damage remain incompletely defined. Here, we investigated cerebral arterial function in mice heterozygous for the *Col4a2^+/G646D^* mutation. Cerebral pial arteries from aged (12-month-old), but not young (3-month-old), *Col4a2^+/G646D^*mice exhibited impaired pressure-induced myogenic tone, whereas downstream parenchymal arterioles retained normal myogenic responsiveness, revealing a segment-selective vascular defect. The frequency of subcellular Ca^2+^ signals (Ca^2+^ sparks) generated by the release of Ca^2+^ through ryanodine receptors (RyR2s) on the sarcoplasmic reticulum (SR) was increased, and large-conductance Ca^2+^-activated K^+^ (BK) channel activity was elevated, in cerebral pial artery smooth muscle cells (SMCs) from aged *Col4a2^+/G646D^*mice. SR Ca^2+^ store load and *Ryr2* transcript abundance were unchanged in these cells, but Stimulated Emission Depletion super-resolution microscopy revealed that RyR2 protein clusters were enlarged compared with controls. Protein kinase A (PKA) activation reproduced RyR2 cluster enlargement, whereas PKA inhibition normalized RyR2 cluster organization and BK channel activity in SMCs from aged *Col4a2^+/G646D^* mice. These findings identify age-associated, PKA-dependent, nanoscale remodeling of RyR2 clusters as a previously unknown mechanism underlying cerebral arterial dysfunction in COL4A2-related disease.

## Introduction

Cerebral small vessel disease (cSVD) is a leading cause of stroke, cognitive impairment, and vascular dementia, yet the molecular mechanisms that disrupt blood vessel structure and function remain incompletely understood (1–4). Human genetic studies implicate basement membrane defects as a causal pathway in cSVD. Among these defects, pathogenic variants of *COL4A1* and *COL4A2*, which encode the α1 and α2 chains of type IV collagen, respectively, cause COL4A1/A2-related disorder, also known as Gould syndrome, an autosomal-dominant and clinically heterogeneous multisystem disorder characterized by neurologic, ocular, renal, muscular, and cerebrovascular manifestations (5–11). The α1 and α2 chains assemble into α1α1α2(IV) heterotrimers, which are principal structural components of cerebral vascular basement membranes. Most pathogenic *COL4A1* and *COL4A2* variants substitute highly conserved glycine residues within the triple-helical domain, leading to impaired collagen folding, secretion, matrix assembly, basement membrane stability, and vascular cell– matrix interactions (12). *COL4A1/A2*-related disease overlaps extensively with sporadic cSVD, presenting as white matter hyperintensities, enlarged perivascular spaces, lacunar infarcts, cerebral microbleeds, and intracerebral hemorrhage (ICH) (13, 14). Consistent with this convergence, genome-wide association studies have linked common variants at the *COL4A1/COL4A2* locus, particularly *COL4A2*, to lacunar ischemic stroke and deep ICH (15, 16). Studies of murine *Col4a1* allelic series demonstrate mutation-position–dependent heterogeneity in cerebrovascular dysfunction and brain pathology, suggesting that distinct *Col4a1* mutations perturb vascular function through different mechanisms (17–19). Although COL4A1 and COL4A2 mutations are established genetic causes of cSVD, the vascular signaling defects that precede ICH, particularly in COL4A2-related disease, remain poorly understood.

Previous studies from our group identified age-dependent ion channel signaling defects in cerebral arterial smooth muscle cells (SMCs) from *Col4a1* mutant mice that impair blood flow and promote ICH (17, 18). In resistance vessels, increases in intraluminal pressure normally evoke the vascular myogenic response, a vital autoregulatory mechanism in which SMC depolarization and constriction stabilize cerebral blood flow and protect downstream capillaries from damaging pressure fluctuations (20). We found that myogenic constriction is impaired in *Col4a1* mutant mice through disruption of TRPM4 (transient receptor potential melastatin 4), a Ca^2+^-activated, monovalent-cation–selective channel required for pressure-induced SMC depolarization, vasoconstriction, and cerebral blood flow autoregulation *in vivo* (17, 18, 21–23). In mice heterozygous for the Gould syndrome-causing *Col4a1* mutation, G394V (*Col4a1*^+/G394V^ mice), age-dependent loss of myogenic tone is caused by depletion of phosphatidylinositol 4,5-bisphosphate [PI(4,5)_2_], an essential cofactor for TRPM4 activation (17). In contrast, *Col4a1*^+/G1344D^ mice, heterozygous for a Gould syndrome-causing *Col4a1* mutation with overlapping symptoms, exhibit a disruption in intracellular Ca^2+^ signaling that impairs TRPM4 activity and myogenic constriction (18). Thus, distinct *Col4a1* mutations converge on impaired pressure-induced autoregulation through divergent signaling mechanisms. These findings raise the possibility that *Col4a2* mutations may also disrupt cerebrovascular function through distinctive mechanistic pathways.

*Col4a2*^+/G646D^ mice develop spontaneous ICH, but the vascular defects that precede hemorrhage have remained unknown (24, 25). Here, we found that cerebral pial arteries from *Col4a2^+/G646D^* mice exhibit age-dependent failure of myogenic tone, similar to that observed in *Col4a1* mutant mice. However, in contrast to *Col4a1* mutants, TRPM4 channel activity was preserved in SMCs from aged *Col4a2^+/G646D^* mice. Instead, myogenic dysfunction was associated with increased activity of large-conductance Ca^2+^-activated K^+^ channels (BK channels). In SMCs, BK channels on the plasma membrane are driven by subcellular Ca^2+^ signaling events (“Ca^2+^ sparks”) generated by Ca^2+^ released from the sarcoplasmic reticulum (SR) through type 2 ryanodine receptors (RyR2s). Super-resolution imaging revealed that RyR2 protein clusters in SMCs from aged *Col4a2^+/G646D^* mice were enlarged compared with controls, and inhibition of protein kinase A (PKA) normalized RyR2 cluster organization and BK channel activity. These findings identify age-dependent, PKA-mediated nanoscale remodeling of RyR2 clusters as a central mechanism of cerebrovascular dysfunction in cSVDs associated with mutant COL4A2.

## Results

### Age-dependent impairment of pressure-induced myogenic tone in cerebral pial arteries, but not parenchymal arterioles, from *Col4a2^+/G646D^* mice

Myogenic tone is impaired in multiple forms of cSVD (17, 26), including those caused by *Col4a1* mutations (17, 18). We therefore used e*x vivo* pressure myography (27) to determine whether the *Col4a2^+/G646D^* mutation (Figure 1A) alters pressure-induced constriction in cerebral resistance vessels (Figure 1 B). Intraluminal pressure was increased stepwise from 5 to 140 mmHg in isolated, cannulated cerebral arteries superfused with Ca^2+^-containing physiological saline, and luminal diameter was recorded once tone stabilized at each step. Passive diameter was then obtained by superfusing vessels with Ca^2+^-free solution, and myogenic tone was calculated as percent constriction normalized to passive diameter.

**Figure 1.**
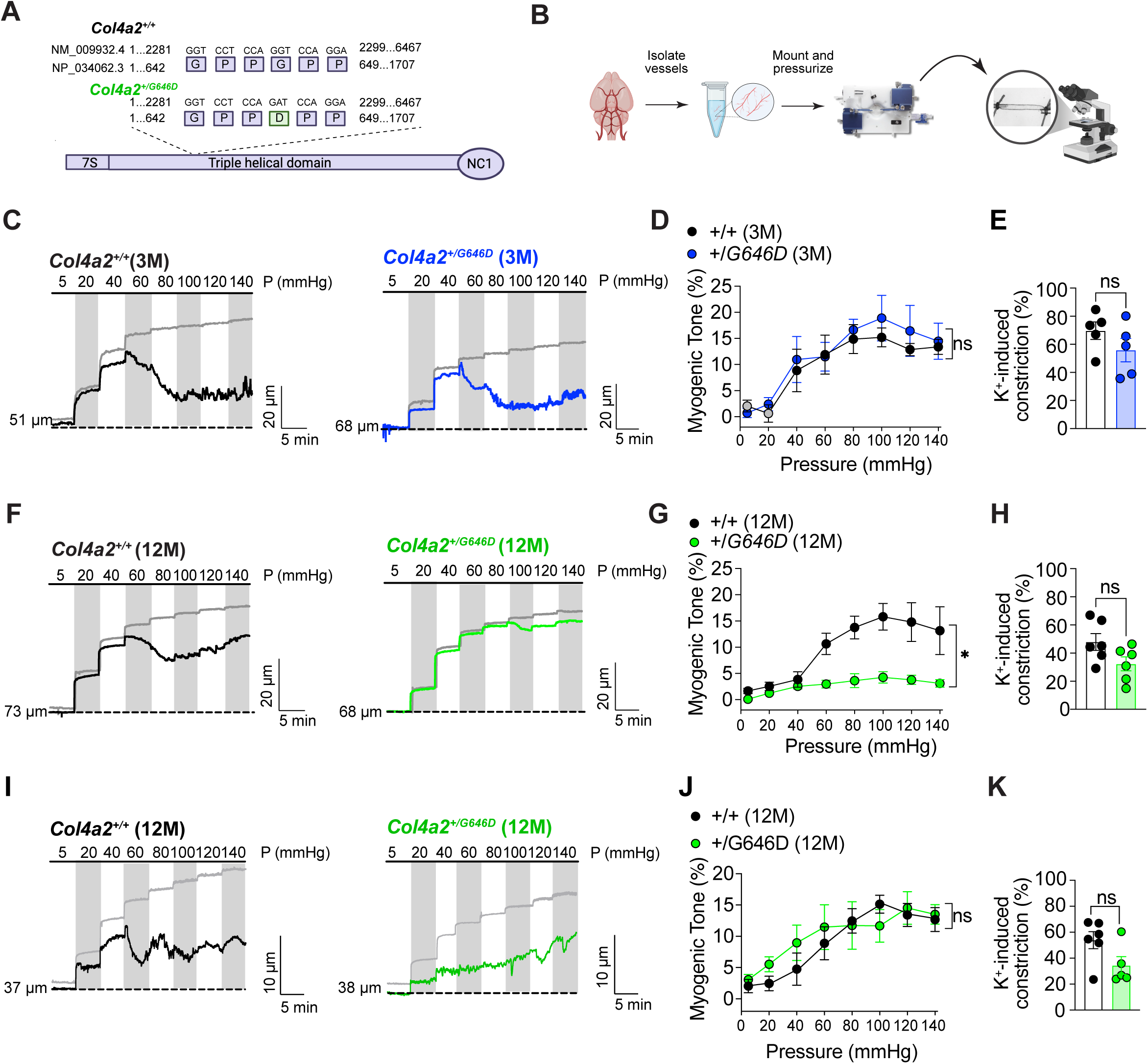
Age-dependent impairment of pressure-induced myogenic tone in cerebral pial arteries, but not parenchymal arterioles, from *Col4a2^+/G646D^* mice. (**A)**, Schematic showing the *Col4a2^+/G646D^* mutation. (**B**), Diagram of *ex vivo* pressure myography experiments. (**C**), Representative pressure-myography recordings from cerebral arteries isolated from 3-month-old *Col4a2*^+/+^ and *Col4a2^+/G646D^* mice during stepwise increases in intraluminal pressure from 5 to 140 mmHg. Active diameter traces are shown in black for arteries from *Col4a2*^+*/+*^ mice and blue for arteries from *Col4a2^+/G646D^* animals; gray traces indicate passive diameter responses. (**D**), Summary plot of myogenic tone as a function of intraluminal pressure in arteries from 3-month-old mice (n = 5 arteries from 3 to 4 animals per group). (**E**), Summary of K^+^-induced constriction of arteries from 3-month-old mice (n = 5 arteries from 3 to 4 animals per group). (**F**), Representative pressure-myography recordings from cerebral arteries isolated from 12-month-old *Col4a2^+/+^* and *Col4a2^+/G646D^* mice. Active diameter traces are shown in black for arteries from *Col4a2^+/+^* mice and green for arteries from *Col4a2^+/G646D^* mice; gray traces indicate passive diameter responses. (**G**), Summary plot of myogenic tone as a function of intraluminal pressure in arteries from 12-month-old mice (n = 6 arteries from 4 animals per group). (**H**), Summary of K^+^-induced constriction of arteries from 12-month-old mice (n = 6 arteries from 4 animals per group). (**I**), Representative pressure-myography recordings from parenchymal arterioles isolated from 12-month-old *Col4a2^+/+^* and *Col4a2^+/G646D^* mice. (**J**), Summary plot of myogenic tone as a function of intraluminal pressure for parenchymal arterioles from 12-month-old mice (n = 5 or 6 arterioles from 3 or 4 animals per group). (**K**), Summary of K^+^-induced vasoconstriction (n = 5 or 6 arterioles from 3 or 4 animals per group). Data are presented as means ± SEM (\**P* <0.05; ns, not significant; 2-way ANOVA with Šídák’s multiple comparisons or unpaired t-test, as appropriate).

Cerebral pial arteries from 3-month-old *Col4a2^+/G646D^*mice developed myogenic tone comparable to that of age-matched *Col4a2*^+/+^ controls (Figure 1, C and D). Constriction in response to 60 mM extracellular K^+^, which directly depolarizes SMCs, was also similar between groups (Figure 1E). In contrast, cerebral pial arteries from 12-month-old *Col4a2^+/G646D^*mice constricted only modestly in response to increasing intraluminal pressure, developing significantly less myogenic tone than age-matched controls (Figure 1, F and G). K^+^-induced constriction remained intact in arteries from 12-month-old *Col4a2^+/G646D^*mice, indicating that the mutation does not cause a generalized impairment of SMC contraction or gross disruption of the contractile apparatus (Figure 1H).

In *Col4a1* mutant mice, myogenic tone is defective in both cerebral pial arteries (17, 18) and parenchymal arterioles (Supplemental Figure 1, A to D). In contrast, cerebral parenchymal arterioles from 12-month-old *Col4a2^+/G646D^* mice developed myogenic tone comparable to that of age-matched *Col4a2*^+/+^ controls, and K^+^-induced constriction was also similar between groups (Figure 1, I to K). These findings identify an age-dependent, segment-selective defect in pressure-induced constriction that is not explained by loss of contractile responsiveness to depolarization in *Col4a2^+/G646D^*mice.

To investigate how segmental loss of myogenic tone alters downstream pressure transmission and flow, we used a simplified series-resistance computational model of the cerebral microcirculation in which pial arteries supply parenchymal arterioles, which in turn supply the arteriole-to-capillary transition (ACT) zone and the capillary bed. Arterial pressure was varied from 70 to 140 mmHg, and simulations were performed with myogenic tone intact, selectively lost in pial arteries, selectively lost in parenchymal arterioles, or lost in both upstream segments. Under intact conditions, capillary inlet pressure and relative capillary flow increased gradually as arterial pressure was raised, consistent with preservation of autoregulatory reserve across the modeled pressure range (Supplemental Figure 2, A and B). In contrast, segmental loss of myogenic tone amplified downstream pressure transmission and flow, with the effects becoming most apparent at pressures above ∼120 mmHg, where autoregulatory reserve was reduced. Loss of parenchymal arteriole tone produced larger increases in capillary inlet pressure and relative capillary flow than loss of pial artery tone alone, consistent with the strong influence of distal resistance on capillary hemodynamics. Notably, combined loss of pial artery and parenchymal arteriole tone produced effects that were greater than the sum of the individual perturbations, indicating synergistic interactions between upstream resistance segments. Thus, in this series-resistance network, loss of myogenic tone in one segment shifts the operating conditions of the remaining segments, amplifying pressure transmission and flow when autoregulatory reserve is challenged.

### BK channel activity is elevated in SMCs from aged *Col4a2^+/G646D^* mice

To determine whether ion channel dysregulation accounts for age-dependent loss of myogenic tone, we first assessed Ca^2+^-dependent activation of TRPM4 using the conventional whole-cell patch-clamp electrophysiology configuration. SMC from 12-month old mice were dialyzed with a high free [Ca^2+^] solution, and cation currents were recorded during application of voltage ramps (−100 to +100 mV). The TRPM4 component of this current was isolated by current subtraction following application of the selective TRPM4 inhibitor NBA (naphthyloxybenzoic acid) (28). We found that Ca^2+^-activated whole-cell TRPM4 currents did not differ between 12-month-old *Col4a2^+/G646D^*mice and age-matched control littermates (Figure 2, A and B).

**Figure 2.**
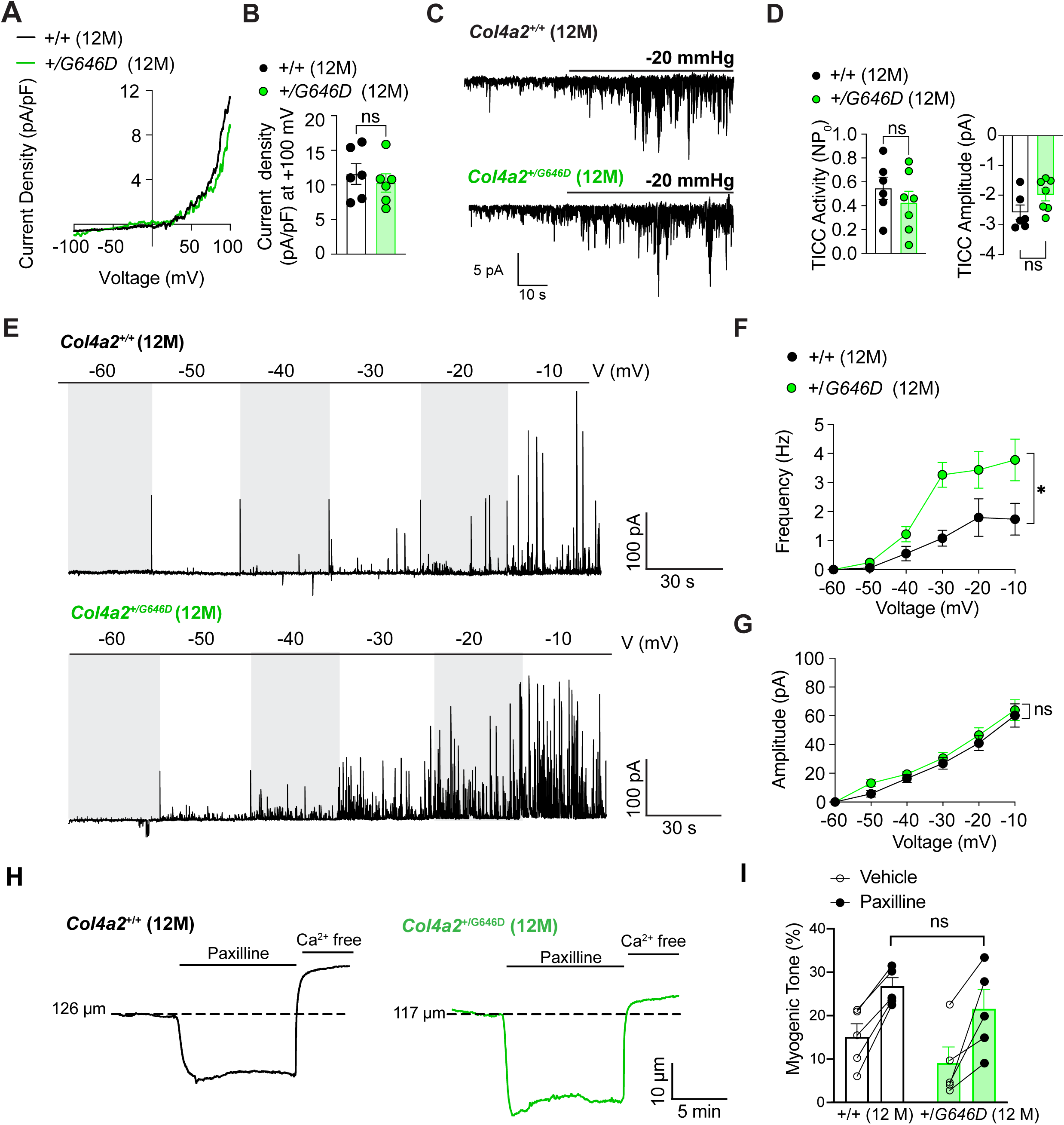
TRPM4 activity is preserved whereas BK channel activity is elevated in SMCs from aged *Col4a2^+/G646D^* mice. (**A)** Representative current-voltage (I-V) relationship of whole-cell TRPM4 currents recorded from cerebral artery SMCs isolated from 12-month-old *Col4a2^+/+^* and *Col4a2^+/G646D^* mice. (**B**) Summary of TRPM4 current density measured at +100 mV (n = 6 cells from 4 animals per group). (**C**) Representative recordings of TICCs in cerebral artery SMCs isolated from 12-month-old *Col4a2^+/+^* and *Col4a2^+/G646D^* mice. TICCs were induced by -20 mmHg and recorded at a holding potential of -70 mV. (**D**) Summary data showing TICC activity (NP_o_) and TICC amplitude (n = 6 or 7 cells from 4 or 5 animals per group). (**E**) Representative recordings of STOCs obtained from SMCs voltage-clamped at membrane potentials ranging from -60 to -10 mV. (**F**) Summary of STOC frequency as a function of membrane potential. (**G**) Summary of STOC amplitude as a function of membrane potential (n = 7 or 9 cells from 3 or 4 animals per group). (**H**) Representative diameter recordin*gs* of pressurized (80 mmHg) cerebral pial arteries isolated from 12-month-old *Col4a2^+/+^* (black) and *Col4a2^+/G646D^* (green) mice before and after superfusion with the selective BK channel blocker paxilline (1 µM), followed by superfusion with Ca^2+^-free PSS to obtain passive diameter. (**I**) Summary data showing myogenic tone (%) before (vehicle) and after paxilline treatment. Open circles, vehicle; filled circles, paxilline; connecting lines link paired measurements from the same vessel (n = 5 vessels from 5 animals per group). Data are presented as means ± SEM (\**P* <0.05; ns, not significant; 2-way ANOVA with Šídák’s multiple comparisons, as appropriate).

We also investigated TRPM4 channel activation in response to mechanical stimulation. Using an established protocol (29), we patch-clamped native cerebral artery SMCs in the amphotericin B-perforated patch configuration and applied suction (−20 mmHg) via the patch pipette to stretch the plasma membrane and stimulate cation currents (22). We found that TRPM4-dependent, stretch-induced transient inward cation currents (TICCs) in SMCs from 12-month-old *Col4a2^+/+^* and *Col4a2^+/G646D^* mice did not differ (Figure 2, C and D). These data indicate that TRPM4 channel activity and associated mechanotransduction pathways are not impaired in SMCs from 12-month-old *Col4a2^+/G646D^*mice.

In SMCs, Ca^2+^ sparks generated by SR Ca^2+^ release through RyR2s activate nearby BK channel clusters in the plasma membrane, producing spontaneous transient outward currents (STOCs). STOCs provide negative feedback hyperpolarization that opposes TRPM4-dependent depolarization, stabilizes the membrane potential, and limits pressure-induced constriction (30). Accordingly, we tested whether enhanced BK channel activity could account for impaired myogenic constriction by assessing STOCs. STOC activity in native cerebral artery SMCs was measured across command potentials from -60 to 0 mV (30, 31) by patch clamping using the amphotericin B– perforated patch configuration. We found that STOC frequency was significantly higher in SMCs from 12-month-old *Col4a2^+/G646D^*mice compared with controls, but amplitude did not differ between groups (Figure 2, E and F). To determine if this effect was age-dependent, we also measured STOCs in cerebral artery SMCs from 3-month-old mice. We found that STOC frequency and amplitude in SMCs from 3-month-old *Col4a2^+/G646D^* mice and age-matched controls did not differ (Supplementary Figure 3, A to C).

To link increased STOC activity with loss of myogenic tone in *Col4a2^+/G646D^* mice, we pressurized isolated cerebral pial arteries to 80 mmHg, allowed them to develop steady-state myogenic tone, and then acutely administered the selective BK channel blocker paxilline. Paxilline increased myogenic tone in cerebral pial arteries from both 12-month-old *Col4a2*^+/+^ and *Col4a2^+/G646D^* mice (Figure 2, H and I). However, following BK channel inhibition, myogenic tone was no longer different between genotypes, suggesting that excessive BK channel activity is the proximate cause of impaired myogenic tone in aged *Col4a2^+/G646D^*arteries.

### Ca^2+^ spark activity is elevated in cerebral artery SMCs from 12-month-old *Col4a2^+/G646D^*mice

Simultaneous intracellular Ca^2+^ and whole cell potassium current recordings indicate that each Ca^2+^ spark generates a STOC (32). We therefore tested whether increased Ca^2+^ spark activity accounts for the elevated STOC frequency observed in SMCs from aged *Col4a2^+/G646D^*mice. Freshly isolated cerebral artery SMCs were loaded with the Ca^2+^-sensitive fluorophore Fluo-4 AM and imaged using high-speed, high-resolution spinning-disk confocal microscopy. Ca^2+^ spark frequency and spatial spread were significantly increased in SMCs from 12-month-old *Col4a2^+/G646D^* mice compared with age-matched *Col4a2^+/+^* controls (Figure 3, A to C; Supplemental Videos 1 and 2). In contrast, Ca^2+^ spark amplitude, duration, rise time, and decay kinetics did not differ between groups (Figure 3, A to C). These findings suggest that elevated STOC frequency in SMCs from *Col4a2^+/G646D^* mice is driven by increased Ca^2+^ spark frequency.

**Figure 3.**
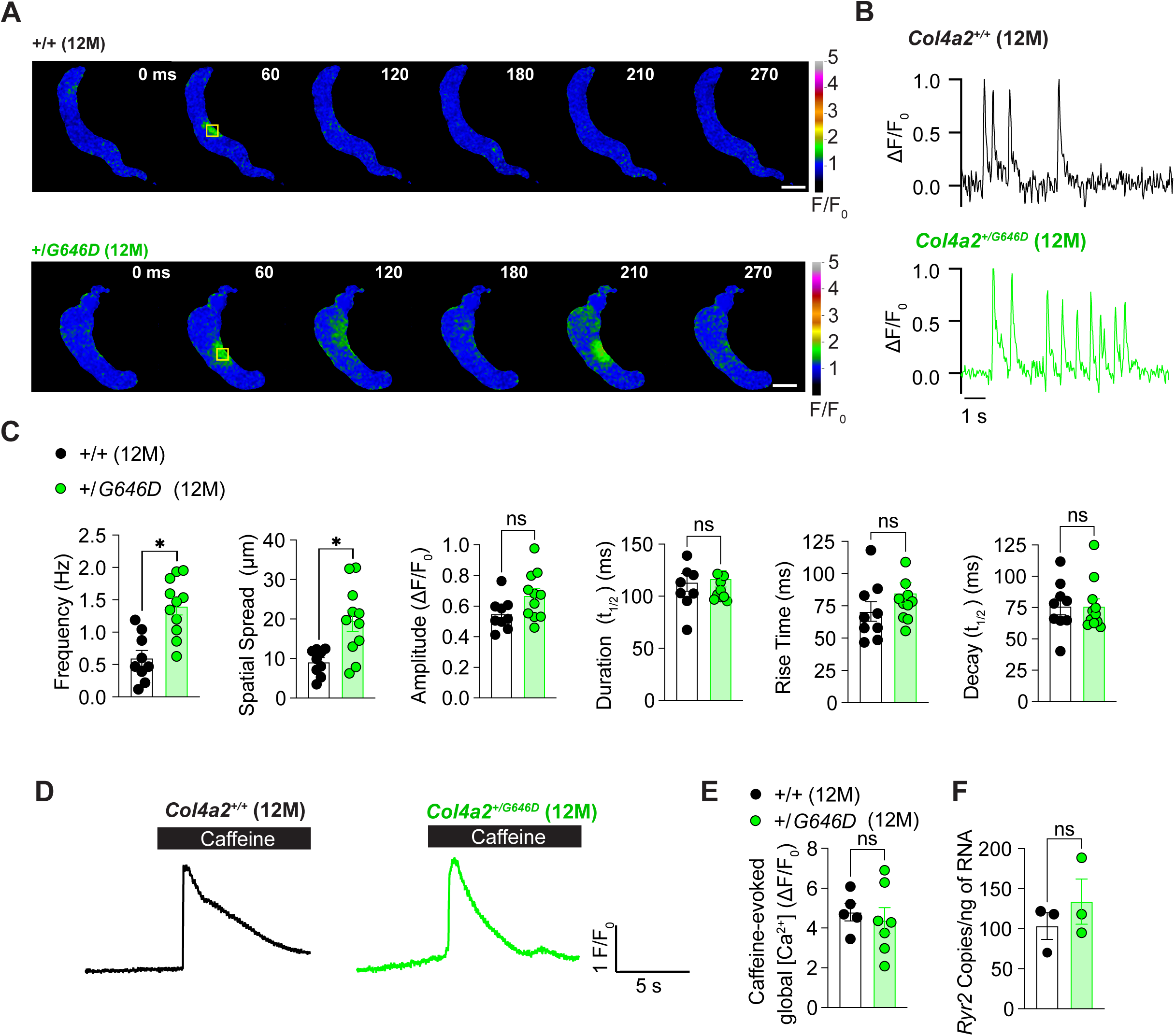
Ca^2+^ spark activity is elevated in cerebral artery SMCs from 12-month-old *Col4a2^+/G646D^* mice. (**A)** Representative time series of Fluo-4 fluorescence images showing spontaneous Ca^2+^ sparks in freshly isolated cerebral artery SMCs from 12-month-old *Col4a2^+/+^* and *Col4a2^+/G646D^* mice. Images are displayed as F/F_0_ at the indicated time points. The yellow boxes indicate the ROIs in which Ca^2+^ sparks were measured. Scale bars: 5 μm. (**B**) Representative ΔF/F_0_ traces from the cells shown in A. Arrowheads indicate individual Ca^2+^ sparks. (**C**) Summary data for Ca^2+^ spark frequency, spatial spread, amplitude, duration, rise time, and decay time in SMCs from 12-month-old *Col4a2^+/+^* and *Col4a2^+/G646D^* mice (n = 8 to 11 cells from 4 animals per group). (**D**) Representative caffeine-evoked global Ca^2+^ transients in Fluo-4-loaded cerebral artery SMCs from 12-month-old *Col4a2^+/+^* and *Col4a2^+/G646D^* mice. Caffeine (10 mM) was applied during the period indicated by the black bars. (**E**) Summary data of caffeine-evoked global Ca^2+^ responses (n = 5 to 7 cells from 4 animals per group). (**F**) ddPCR quantification of *Ryr2* transcript abundance in cerebral arteries from 12-month-old *Col4a2^+/+^* and *Col4a2^+/G646D^* mice (n = 3 mice per group). Data are presented as means ± SEM (\**P* <0.05; ns, not significant; unpaired t-test).

Increased SR Ca^2+^ load can increase Ca^2+^ spark frequency (33, 34). Therefore, we compared the relative SR Ca^2+^ content between groups by exposing Fluo-4 AM-loaded SMCs to caffeine (5 mM) to maximally stimulate RyR2-mediated SR Ca^2+^ release into the cytosol. Caffeine-evoked Ca^2+^ transients did not differ between SMCs from 12-month-old *Col4a2^+/+^* and *Col4a2^+/G646D^* mice (Figure 3, D and E), suggesting that increased Ca^2+^ spark activity is not driven by elevated SR Ca^2+^ load.

We also tested whether increased *Ryr2* expression could account for enhanced Ca^2+^ spark activity. Quantitative droplet digital PCR (ddPCR) revealed no difference in *Ryr2* transcript abundance between SMCs from 12-month-old *Col4a2^+/+^* and *Col4a2^+/G646D^*mice (Figure 3F). Together, these data indicate that *Col4a2^+/G646D^*SMCs exhibit increased Ca^2+^ spark activity without detectable increases in SR Ca^2+^ content or *Ryr2* mRNA abundance.

### Nanoscale remodeling of RyR2 protein clusters in cerebral artery SMCs from *Col4a2^+/G646D^* mice

RyR2s assemble into multichannel clusters within the SR membrane (35). Computational and experimental studies indicate that the size and spatial arrangement of RyR2 clusters influence the probability of Ca^2+^ release, with larger clusters associated with higher Ca^2+^ spark frequency (36–39). Accordingly, we investigated whether the nanoscale organization of RyR2 clusters was altered in SMCs from 12-month-old *Col4a2^+/G646D^* mice. Freshly-isolated SMCs were fixed, immunolabeled for RyR2 (40), and imaged using Stimulated Emission Depletion (STED) super-resolution microscopy (41). Prominent RyR2 clusters were present in both groups; however, clusters were significantly larger in SMCs from 12-month-old *Col4a2^+/G646D^* mice than in controls. Based on an estimated cytoplasmic RyR2 footprint of ∼900 nm^2^ per channel (35), and assuming close packing within each cluster, we estimate that mean-sized RyR2 clusters contain approximately 10 channels in SMCs from controls and approximately 15 channels in SMCs from *Col4a2^+/G646D^* mice (Figure 4, A and B). In contrast, RyR2 cluster density was lower in SMCs from *Col4a2^+/G646D^* mice compared with age-matched controls (Figure 4, A and B). Thus, the *Col4a2^+/G646D^* mutation is associated with redistribution of RyR2 into fewer, larger clusters, a pattern consistent with coalescence or reorganization of preexisting RyR2 clusters rather than increased RyR2 expression.

**Figure 4.**
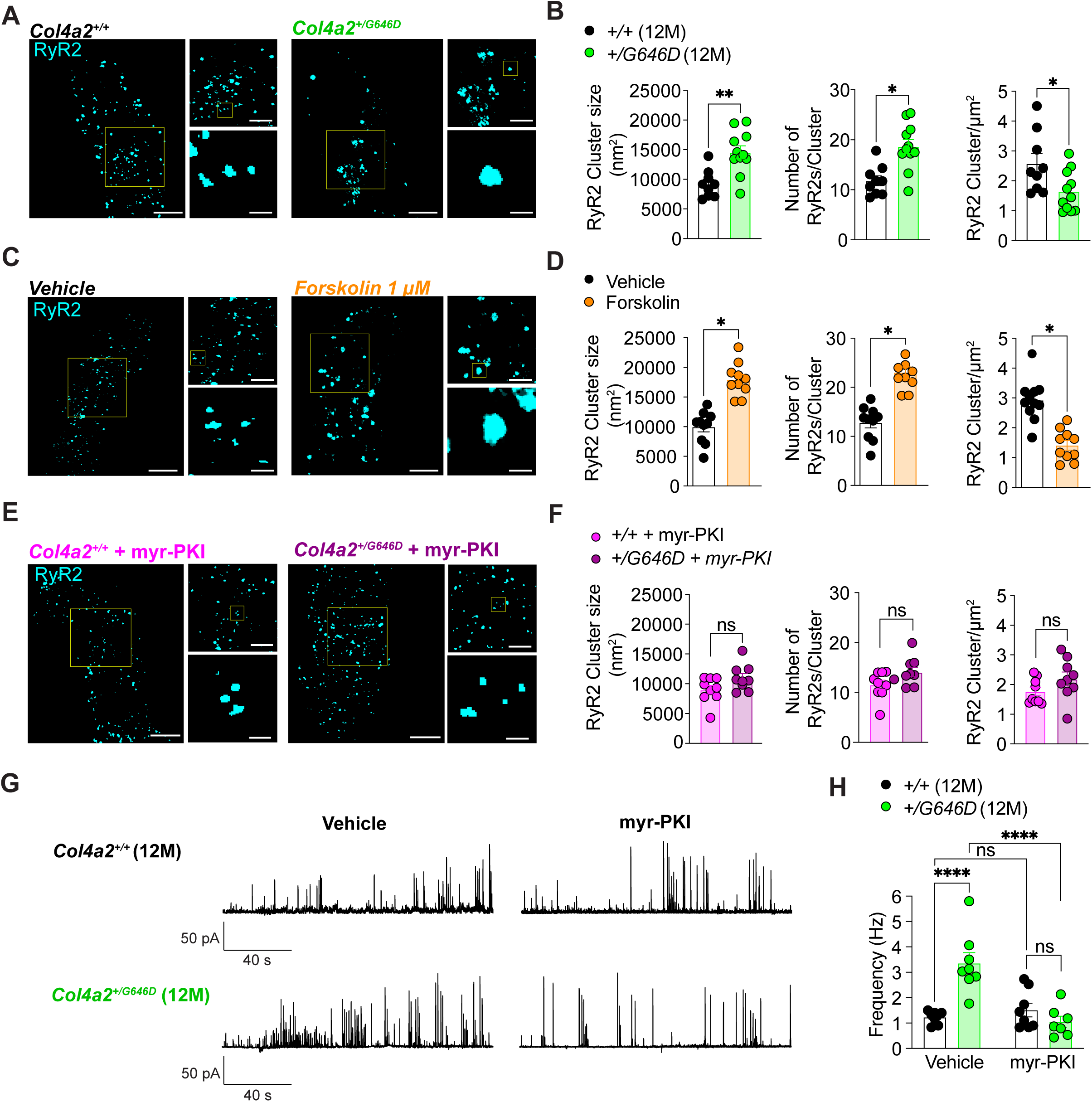
PKA-dependent remodeling of RyR2 protein clusters in cerebral artery SMCs from aged *Col4a2^+/G646D^* mice. (**A)** Representative STED super-resolution images of RyR2 immunolabeling in freshly isolated cerebral artery SMCs from 12-month-old *Col4a2^+/+^* and *Col4a2^+/G646D^* mice. Yellow boxes indicate regions shown at higher magnification. (**B**) Summary data showing RyR2 cluster size, estimated number of RyR2 channels per cluster, and RyR2 cluster density in SMCs from 12-month-old *Col4a2^+/+^* and *Col4a2^+/G646D^* mice (n = 9 to 11 cells from 4 animals per group). (**C**) Representative STED images of RyR2 immunolabeling in cerebral artery SMCs treated with vehicle or forskolin (1 μM). Yellow boxes indicate magnified regions. (**D**) Summary data showing the effects of forskolin on RyR2 cluster size, estimated RyR2 channel number per cluster, and RyR2 cluster density (n = 9 to 10 cells from 4 animals per group). (**E**) Representative STED images of RyR2 immunolabeling in cerebral artery SMCs from 12-month-old *Col4a2^+/+^* and *Col4a2^+/G646D^* mice treated with the PKA inhibitor myr-PKI (300 nM). Yellow boxes indicate magnified regions. (**F**) Summary data showing RyR2 cluster size, estimated RyR2 channel number per cluster, and RyR2 cluster density after myr-PKI treatment (n = 9 cells from 4 animals per group). Scale bars = 3 µm (left), 1 µm (top right), and 200 nm (bottom right). (**G**) Representative recordings of STOCs using the perforated-patch configuration in cerebral artery SMCs from 12-month-old *Col4a2^+/+^* and *Col4a2^+/G646D^* mice treated with the PKA inhibitor myr-PKI (300 nM). (**H**) Summary data of STOC frequency from cells treated with vehicle or myr-PKI (n = 8 cells from 5 animals per group). Data are presented as means ± SEM (\**P* <0.05; \*\**P* <0.01; \*\*\*\**P* <0.0001; ns, not significant; unpaired t-test or 2-way ANOVA with Šídák’s multiple comparisons, as appropriate). Each symbol represents one cell. (**I**) Proposed mechanism in which PKA-dependent nanoscale remodeling of RyR2 clusters increases Ca^2+^ spark and STOC frequency to oppose pressure-induced vasoconstriction in cerebral arteries from *Col4a2^+/G646D^* mice.

Previous studies reported that generation of cyclic adenosine monophosphate (cAMP) by adenylyl cyclase and subsequent activation of PKA increases Ca^2+^ spark frequency in SMCs and cardiomyocytes (42–44), and that RyR2 channel activity is regulated by PKA-mediated phosphorylation (45, 46). In addition, computational modeling predicts that PKA-mediated phosphorylation can promote RyR2 cluster reorganization (42). We therefore tested whether activation of the adenylyl cyclase/cAMP/PKA pathway is sufficient to remodel RyR2 clusters in cerebral artery SMCs by treating freshly isolated SMCs from 3-month-old C57BL/6J mice with the adenylyl cyclase activator forskolin (1 μM). Consistent with previous work showing increased STOC frequency after forskolin treatment (44), it also increased RyR2 cluster size and reduced cluster density, phenocopying the RyR2 cluster organization observed in SMCs from *Col4a2^+/G646D^* mice (Figure 4, C and D). These data indicate that activation of the adenylyl cyclase signaling cascade is sufficient to drive RyR2 cluster remodeling in cerebral artery SMCs.

We next assessed whether PKA activity is responsible for enlarged RyR2 clusters in SMCs from 12-month-old *Col4a2^+/G646D^* mice by treating isolated SMCs with the selective PKA inhibitor myr-PKI (300 nM, 10 min) before fixation and RyR2 immunolabeling (47). Following myr-PKI treatment, RyR2 cluster size and density no longer differed between SMCs from *Col4a2^+/G646D^* mice and age-matched controls (Figure 4, E and F), indicating that sustained PKA activity underlies RyR2 cluster remodeling in SMCs from aged *Col4a2^+/G646D^* mice.

### PKA inhibition restores STOC frequency in cerebral artery SMCs from *Col4a2^+/G646D^* mice

Because PKA inhibition restored the normal distribution of RyR2 clusters, we tested whether this change was also accompanied by a reduction in STOC frequency. The frequency of STOCs in voltage-clamped (−30 mV) SMCs from 12-month-old *Col4a2^+/G646D^* mice treated with myr-PKI did not differ from that in age-matched controls (Figure 4, G and H). Together, these results support a model in which PKA-dependent nanoscale remodeling of RyR2 clusters increases Ca^2+^ spark and STOC frequency to oppose pressure-induced vasoconstriction in cerebral arteries from *Col4a2^+/G646D^*mice (Figure 4I).

## Discussion

This study identifies a previously unrecognized mechanism of cerebral artery dysfunction in COL4A2-related disease. We found that cerebral pial arteries from aged *Col4a2^+/G646D^* mice exhibit selective, age-dependent loss of myogenic tone, whereas parenchymal arterioles do not. Myogenic dysfunction was associated with increased Ca^2+^ spark frequency, elevated BK channel activity, and nanoscale remodeling of RyR2 protein clusters into larger signaling domains. Pharmacological activation of adenylyl cyclase in SMCs from wild-type mice reproduced this RyR2 cluster phenotype, whereas inhibition of PKA normalized RyR2 cluster organization and BK channel activity in SMCs from *Col4a2^+/G646D^*mice. Together, these findings support a model in which age-dependent activation of PKA in *Col4a2* mutants remodels the RyR2 signaling architecture, increasing the frequency of localized SR Ca^2+^-release events and BK channel-dependent hyperpolarization that oppose pressure-induced smooth muscle depolarization and constriction. This mechanism distinguishes *Col4a2^+/G646D^*mice from previously characterized *Col4a1* mutant models and provides further evidence that COL4A1/A2-related cerebrovascular diseases arise through mutation-specific disruption of distinct ion channel signaling networks.

Preservation of myogenic tone in parenchymal arterioles from 12-month-old *Col4a2^+/G646D^* mice distinguishes this model from previously characterized *Col4a1^+/G394V^* and *Col4a1^+/G1344D^*mutants, in which pressure-induced constriction is impaired in both cerebral pial arteries and parenchymal arterioles (17, 18). The segment-selective loss of function in aged *Col4a2^+/G646D^* mice reflects differences in the primary smooth muscle ion channel defects caused by these mutations. In *Col4a1* mutants, impaired TRPM4 activity compromises myogenic tone throughout the cerebral resistance vasculature because TRPM4 channels are required for pressure-induced depolarization in both pial arteries and parenchymal arterioles (21, 22, 48). In contrast, *Col4a2^+/G646D^* mice exhibit hyperactivity of the RyR2/Ca^2+^ spark/BK channel pathway, which is functionally prominent in pial artery SMCs but has limited influence in parenchymal arteriole SMCs, where Ca^2+^ sparks are not generated under normal physiological conditions, and BK channel activity has minimal impact on membrane potential (26).

Additional evidence from *Col4a1* mutant models demonstrates that distinct collagen IV mutations perturb the cerebrovascular network through segment-specific mechanisms. In the retinal vasculature of *Col4a1*^+/G498V^ and *Col4a1*^+/G1064D^ mice, the ACT zone, a region covered by ensheathing pericytes (49), is hypermuscularized, with mural cell hyperplasia and increased contractile protein expression (50). Computational models suggested that ACT zone remodeling increases upstream intravascular pressure in parenchymal arterioles, thereby promoting ICH (50). In *Col4a2^+/G646D^* mice, by contrast, loss of myogenic tone in cerebral pial arteries with preserved tone in downstream parenchymal arterioles is predicted to create a distinct hemodynamic phenotype. A simplified series-resistance model predicted that selective loss of pial artery tone reduces upstream resistance and increases transmission of arterial pressure to the capillary inlet, even when parenchymal arteriole myogenic responsiveness is preserved. However, intact downstream parenchymal arteriole tone limits this effect, indicating that preserved parenchymal arteriole constriction can compensate for loss of upstream pial artery tone over part of the autoregulatory range. This compensation becomes less effective at arterial pressures above ∼120 mmHg, where segmental differences in autoregulatory reserve become more apparent and capillary inlet pressure and flow rise more steeply. The model also predicts that the combined loss of pial artery and parenchymal arteriole tone produces a super-additive increase in downstream pressure transmission and flow. This nonlinear behavior arises naturally from the series-resistance structure of the vascular network, because loss of buffering in one segment changes the pressure-flow operating point of the other. Thus, preserved parenchymal arteriole tone in *Col4a2^+/G646D^* mice might protect the capillary bed from the full consequences of pial artery dysfunction, but at the cost of reduced autoregulatory reserve and increased dependence on downstream microvascular resistance to buffer pressure fluctuations.

Ion channel clustering strongly influences channel regulation, gating behavior, and downstream signaling (51). Incorporation of multiple RyR2 channel monomers into a cluster increases the probability that spontaneous opening of a single unit will recruit neighboring RyR2s through Ca^2+^-induced Ca^2+^ release, thereby increasing the likelihood and spatial spread of local Ca^2+^ spark events (37, 52). RyR2 cluster enlargement has been implicated in pathological Ca^2+^ release in cardiovascular disease (36, 53), including in prior work from our group showing that increased RyR2 cluster size in cerebral artery SMCs enhances Ca^2+^ spark and BK channel activity and disrupts myogenic tone in *mdx* mice (36), a model of Duchenne muscular dystrophy. The present findings extend this concept to COL4A2-related cerebrovascular disease by showing that stimulation of the adenylyl cyclase/cAMP/PKA pathway is sufficient to enlarge RyR2 clusters, and that PKA inhibition normalizes RyR2 cluster size in SMCs from aged *Col4a2^+/G646D^* mice. These results are consistent with previous studies showing that cAMP/PKA-dependent phosphorylation increases RyR2 activity (45, 46, 53), that forskolin increases the frequency of Ca^2+^ sparks in ventricular cardiomyocytes and cerebral SMCs (44, 54), and that PKA activation promotes RyR2 cluster enlargement (55). Together, these findings suggest that PKA-dependent reorganization of RyR2 clusters enhances subcellular Ca^2+^ release, increases BK channel activity, and impairs myogenic tone in cerebral arteries from *Col4a2*^+/*G646D*^ mice.

Two key unresolved questions remain: how does a point mutation in *Col4a2* increase PKA-dependent signaling in SMCs, and why does this process emerge with age? PKA is commonly activated downstream of Gα_s_-coupled GPCRs at the plasma membrane, raising the possibility that mutant collagen IV alters signaling through extracellular matrix-sensitive adhesion G-protein–coupled receptors (GPCRs) (56). GPR126/ADGRG6 is a Gα_s_-coupled adhesion GPCR that stimulates cAMP production in response to collagen IV binding (57, 58). Notably, human genetic studies have linked GPR126/ADGRG6 mutations to lacunar stroke (16), and endothelial-specific deletion of *Adgrg6* impairs cerebrovascular function and blood-brain barrier integrity in mice (59). Thus, one plausible model is that mutation-associated disruption of COL4A2 structure, together with age-dependent turnover, remodeling or disorganization of the cerebral vascular basement membrane, alters GPR126/ADGRG6 engagement or activation, thereby enhancing cAMP/PKA signaling in blood vessels. Whether this pathway is activated directly in SMCs from aged *Col4a2^+/G646D^* mice remains an important question for future studies.

In summary, our findings identify PKA-dependent nanoscale remodeling of RyR2 signaling domains as a previously unrecognized mechanism of cerebral arterial dysfunction in COL4A2-related disease. The *Col4a2^+/G646D^* mutation enhances RyR2/Ca^2+^ spark signaling in cerebral artery SMCs, increasing BK-channel-dependent hyperpolarizing currents that oppose pressure-induced constriction. This mechanism produces segment-selective vascular dysfunction, in which myogenic tone is lost in cerebral pial arteries but preserved in parenchymal arterioles, a pattern predicted to reduce autoregulatory reserve. More broadly, these results support the concept that COL4A1/A2 mutations disrupt cerebrovascular function through mutation-specific effects on distinct ion channel signaling networks. Defining how mutations in basement membrane proteins engage intracellular signaling pathways that remodel ion channel architecture may reveal new therapeutic strategies to preserve regulation of cerebral blood flow and reduce hemorrhagic risk in COL4A1/A2-related cSVD.

## Author contributions

SE and DBG initiated and supervised the project. SE, BL, and EY designed the experiments. DBG and CLD provided mutant mice used for this study. YFE assisted with establishing, maintaining, and managing mouse colonies. BL, EY, and ASS performed and analyzed patch-clamp electrophysiology experiments. EY and WZ performed pressure myography experiments. SE performed computational modeling. BL and EY performed Ca^2+^-imaging experiments. BL performed STED microscopy. BL, EY, WZ, ASS, and SE analyzed the data. BL and SE wrote the initial manuscript draft and prepared the figures. SE, BL, EY, WZ, ASS, CLD, YFE, and DBG revised and approved the manuscript.

## Acknowledgments

Funding for this study was provided by NIH/NHLBI (R35HL155008 to SE, R01HL122770 to YFE), NIH/NINDS (RF1NS148111 to SE and R01NS140370 to DBG), NIH/NIA (R01AG097446 to SE), and NIH/NIDDK (R01DK135621 to YFE). DBG is also supported by UCSF Vision Core (NIH/NEI P30EY037668), and by an unrestricted grant from Research to Prevent Blindness, New York, NY to the UCSF Department of Ophthalmology. STED imaging was performed with support from the URMC Center for Advanced Microscopy and Nanoscopy (RRID:SCR_023177) and funding from NIH/HD and DHHS/PHS/NIH (P50HD103536 to the Intellectual and Developmental Disabilities Research Center; S10OD023440 to the Instrumentation Program Award)

## Materials and Methods

### Sex as a biological variable

Young adult (3-month-old) and aged (12-month-old) male and female littermate *Col4a2^+/+^* and *Col4a2^+/G646D^* mice were used in this study. No sex-related differences in the data presented here were observed.

### Chemicals and reagents

Unless otherwise specified, chemicals and other reagents were obtained from Sigma-Aldrich, Inc.

### Animals

The *Col4a2^G646D^* mutation was backcrossed to C57BL/6J mice for over 20 generations (60, 61). Animals were maintained in individually ventilated cages (≤5 mice/cage) with *ad libitum* access to food and water in a room with controlled 12-hour light and dark cycles. Mice were euthanized by decapitation and exsanguination under isoflurane anesthesia (Baxter Healthcare). Brains were isolated and placed in ice-cold Ca^2+^-free physiological saline solution (Mg-PSS), containing 140 mM NaCl, 5 mM KCl, 2 mM MgCl_2_, 10 mM HEPES, and 10 mM glucose (pH 7.4, NaOH), supplemented with 0.5% bovine serum albumin (BSA).

### Isolation of cerebral arteries and SMCs

Cerebral pial arteries (middle cerebral, posterior cerebral, and superior cerebellar) and parenchymal arterioles branching from middle cerebral arteries were gently removed from the brain and washed in Mg-PSS. Native SMCs for all single-cell experiments were obtained by initially digesting isolated pial arteries in 1 mg/mL papain (Worthington Biochemical Corporation), 1 mg/mL dithiothreitol (DTT), and 10 mg/mL BSA in Mg-PSS at 37°C for 12 minutes, followed by a 14-minute incubation with 1 mg/mL type II collagenase (Worthington Biochemical Corporation). A single-cell suspension was prepared by washing digested arteries three times with Mg-PSS and triturating with a fire-polished glass pipette. All cells used for this study were freshly dissociated on the day of experimentation.

### Pressure myography

The current best practices guidelines for pressure myography experiments were followed (27). Arteries or arterioles were mounted between two glass cannulas (outer diameter, 40–50 μm) in a pressure myograph chamber (Living Systems Instrumentation) and secured by a nylon thread. Intraluminal pressure was controlled using a servo-controlled peristaltic pump (Living Systems Instrumentation). Preparations were visualized with an inverted microscope (Accu-Scope Inc.) coupled to a USB camera (The Imaging Source LLC). Changes in luminal diameter were assessed using IonWizard software (version 7.2.7.138; IonOptix LLC, Westwood, MA, USA). Arteries were bathed in warmed (37°C), oxygenated (21% O_2_, 6% CO_2_, 73% N_2_) PSS (119 mM NaCl, 4.7 mM KCl, 21 mM NaHCO_3_, 1.17 mM MgSO_4_, 1.8 mM CaCl_2_, 1.18 mM KH_2_PO_4_, 5 mM glucose, 0.03 mM EDTA) at an intraluminal pressure of 5 mmHg. Following equilibration for 15 minutes, intraluminal pressure was increased to 110 mmHg, and vessels were stretched to their approximate *in vivo* length, after which pressure was reduced back to 5 mmHg for an additional 15 minutes. Vessel viability was assessed for each preparation by evaluating vasoconstrictor responses to high extracellular [K^+^] PSS, made isotonic by adjusting [NaCl] (60 mM KCl, 63.7 mM NaCl). Arteries that showed less than 10% constriction in response to elevated [K^+^] were considered damaged and were excluded from further investigation.

Myogenic tone was assessed by raising intraluminal pressure in 20-mmHg increments from 5 mmHg to 140 mmHg (5 to 20 mmHg for the first step). The active diameter was obtained by allowing vessels to equilibrate for at least 5 minutes at each pressure or until a steady-state diameter was reached. Following completion of the pressure-response study, intraluminal pressure was lowered to 5 mmHg, and arteries were superfused with Ca^2+^-free PSS supplemented with EGTA (2 mM) and the voltage-dependent Ca^2+^ channel blocker diltiazem (10 μM) to inhibit SMC contraction, after which passive diameter was obtained by repeating the stepwise increase in intraluminal pressure. Myogenic tone at each pressure step was calculated as myogenic tone (%) = [1 – (active lumen diameter/passive lumen diameter)] × 100.

### Computational modeling

A simplified single-path resistance model was used to estimate pressure transmission through the cerebral resistance vasculature. The modeled pathway consisted of one pial artery connected in series to one parenchymal arteriole, one arteriole-to-capillary transition (ACT) segment, and a downstream capillary bed. Pial inlet pressure was varied from 70 to 200 mmHg in 5-mmHg increments, and terminal capillary-bed pressure was fixed at 5 mmHg. Segmental resistance was calculated using a relative Poiseuille relationship:

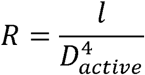

where *l* is the effective segment length and *D_active_* is the active luminal diameter. Active diameter was calculated as:

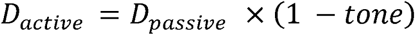

Baseline assumptions were as follows: pial artery passive diameter, 50 μm; pial artery effective length, 3000 μm; baseline pial tone, 15%; maximum pial tone, 30%; parenchymal arteriole passive diameter, 35 μm; parenchymal arteriole effective length, 500 μm; baseline parenchymal arteriole tone, 25%; maximum parenchymal arteriole tone, 40%; ACT passive diameter, 15 μm; ACT effective length, 100 μm; and ACT tone, fixed at 20%. Capillary-bed resistance was calibrated such that capillary inlet pressure was 25 mmHg at 70 mmHg pial inlet pressure under baseline intact-tone conditions.

Pial artery and parenchymal arteriole tone were modeled as pressure-reactive variables that increased linearly with local midpoint pressure until reaching the imposed maximum tone for each segment. In simulations of tone loss, tone in the affected segment was set to zero across all pressures. The ACT segment was treated as a static resistance in all simulations. Total flow was calculated as:

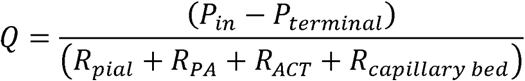

Segmental pressure drops were calculated as Q x R, allowing determination of pressure at the ACT inlet and capillary inlet. Because pressure-reactive tone depends on local midpoint pressure, which in turn depends on flow and segmental pressure drops, solutions were obtained iteratively until the pressures converged. Flow values were normalized to the intact-tone condition at 70 mmHg.

### Electrophysiological recordings

Enzymatically isolated native SMCs were transferred to a recording chamber (Warner Instruments) and allowed to adhere to glass coverslips for 15 minutes at room temperature. Recording electrodes (3–5 MΩ) were pulled and polished. For all patch-clamp electrophysiology experiments, currents were recorded at room temperature using an Axopatch 200B amplifier (Molecular Devices) equipped with an Axon CV 203BU headstage and Digidata 1440A digitizer (Molecular Devices). Whole-cell TRPM4 currents were recorded in a bath solution consisting of 156 mM NaCl, 1.5 mM CaCl_2_, 10 mM glucose, 10 mM HEPES, and 10 mM TEACl (pH 7.4, adjusted with NaOH). The patch-pipette solution contained 156 mM CsCl, 8 mM NaCl, 1 mM MgCl_2_, 10 mM HEPES (pH 7.4, adjusted with NaOH), and a free Ca^2+^ concentration of 200 μM, adjusted using the appropriate amounts of CaCl_2_ and EGTA, as determined with Max-Chelator software (WEBMAXC standard). Whole-cell cation currents were evoked by applying 400-ms voltage ramps from -100 to +100 mV from a holding potential of -60 mV. Voltage ramps were repeated every 2 seconds for 300 seconds. The selective TRPM4 inhibitor, 4-chloro-2-[2-(naphthalene-1-yloxy) acetamido] benzoic acid (NBA; 3 μM), was applied after peak TRPM4 current was recorded (∼100 seconds). Whole-cell TRPM4 current amplitude was expressed as the NBA-sensitive current at +100 mV.

TICCs and STOCs were recorded using the amphotericin B-perforated patch-clamp configuration. The pipette solution contained 110 mM K-aspartate, 1 mM MgCl_2_, 30 mM KCl, 10 mM NaCl, 10 mM HEPES, and 50 μM EGTA at pH 7.2 (KOH). Amphotericin B (200 μg/mL) was included in the pipette solution of all perforated-patch recordings to allow electrical access. Perforation was deemed acceptable if series resistance was less than 50 MΩ. The bathing solution for these recordings was 134 mM NaCl, 6 mM KCl, 1 mM MgCl_2_, 2 mM CaCl_2_, 10 mM HEPES, and 10 mM glucose at pH 7.4 (NaOH). Stretch-induced TICCs were recorded from SMCs voltage-clamped at -70 mV. Membrane stretch was delivered by applying negative pressure (−20 mmHg) through the recording electrode using a pressure clamp system (HSPC-1; ALA Scientific Instruments Inc., Farmingdale, NY, USA). TICC activity was calculated as the sum of the open channel probability (NP_o_) of multiple open states of 1.75 pA (29).

STOCs were recorded from SMCs voltage-clamped over a range of membrane potentials (−60 to -10 mV). STOCs were defined as transient events >10 pA, and the frequency was calculated by dividing the number of events by the time between the first and last event.

### Ca^2+^ imaging

Images were acquired at a rate of 33 frames per second (fps) using an iXon 897 electron-multiplying charge-coupled device (EMCCD) camera (Andor; 16 μm–by–16 μm pixel size) coupled to a spinning-disk confocal head (CSUX1; Yokogawa) with a 100× oil-immersion objective (numerical aperture,1.45; Olympus). For imaging of Ca^2+^ sparks, a suspension of freshly isolated SMCs was placed in a glass-bottom 35-mm dish, and cells were allowed to adhere to the glass coverslip for 20 minutes at room temperature. SMCs were loaded with the Ca^2+^-sensitive fluorophore, Fluo-4 AM (1 μM; Invitrogen), in the dark for 20 minutes at room temperature in Mg^2+^-PSS, washed three times with Ca^2+^-containing PSS, and incubated at room temperature for 20 minutes in the dark to allow sufficient time for Fluo-4 AM de-esterification. Custom software (SparkAn) developed by Dr. A. D. Bonev (University of Vermont) was used to analyze Ca^2+^ spark frequency, amplitude, rise time, decay time, duration, and spatial spread. The threshold for Ca^2+^ spark detection was defined as a local increase in fluorescence ≥ 0.2 ΔF/F_0_.

SR Ca^2+^ loading was imaged by loading cells with Fluo-4 AM as above, then washing three times and imaging in Mg^2+^-PSS to eliminate Ca^2+^-influx signals. Caffeine (10 mM) was applied to the bath to promote release of SR Ca^2+^ through RyR2s.

### Isolation of RNA and ddPCR

Total RNA was extracted from isolated cerebral arteries by homogenization in TRIzol reagent (Invitrogen), followed by purification using a Direct-zol RNA microprep kit (Zymo Research) with on-column DNAse treatment. RNA concentration was determined using an RNA 6000 Pico Kit on a Bioanalyzer 2100 running Agilent 2100 Expert Software (B.02.11; Agilent Technologies). RNA was converted to cDNA using iScript cDNA Supermix (Bio-Rad, Hercules). Quantitative ddPCR was performed using QX200 ddPCR EvaGreen Supermix (Bio-Rad), cDNA templates, and the custom-designed primers for *Ryr2* (NM_023868.2), 5’-TGGTCAGCGTGTCCTCAGAAAGG-3’ (sense) and 5’-CGCCAATGAGATAGCCTTGGGCT-3’ (antisense). Generated droplet emulsions were amplified using a C100 Touch Thermal Cycler (Bio-Rad), and the fluorescence intensity of individual droplets was measured using a QX200 Droplet Reader (Bio-Rad) running QuantaSoft (version 1.7.4; Bio-Rad). Analyses were performed using QuantaSoft Analysis Pro (version 1.0596; Bio-Rad).

### Immunofluorescence

Freshly isolated SMCs were seeded onto a 12-mm coverslip coated with poly-L-lysine for 20 minutes. Cells were fixed with 4% paraformaldehyde in PBS (pH 7.4), for 15 minutes at 4°C, followed by three wash steps for 10 minutes each with PBS. Cells were blocked and permeabilized with 50% Seablock and 0.1% Triton X-100 for 2 hours at room temperature with shaking. Primary antibody incubation was performed overnight with a mouse anti-RyR2 antibody (C3-33; Thermo Fisher), diluted 1:50 in an antibody diluent solution containing 20% Seablock, 0.05% Triton X-100, and 1% BSA. Coverslips were then washed three times for 10 minutes each with antibody diluent solution and incubated for 2 hours with abberior STAR RED-conjugated goat anti-mouse IgG (abberior, #STRED), diluted 1:200 in antibody diluent solution. Coverslips were washed three times with PBS for 10 minutes each at room temperature and mounted with ProLong Gold Antifade (Thermo Fisher, #P36930).

### Stimulated Emission Depletion (STED) microscopy

RyR2 protein clusters in isolated SMCs were imaged using STED microscopy at the Center for Advanced Light Microscopy and Nanoscopy, University of Rochester Medical Center. Images were acquired on an Abberior easy3D STED microscope built around an inverted Olympus IX83 microscope equipped with a 100x/1.40 NA oil-immersion objective. Excitation was provided by a 640-nm diode laser, with stimulated emission depletion performed using a 775-nm depletion laser. Fluorescence emission was detected using four spectral avalanche photodiode detectors. Samples were visualized using Abberior Lightbox iMSPECTOR software (version 16.3). Images were analyzed using ImageJ/Fiji.

### Statistical analysis

Summary data are presented as means ± SEM. Statistical analyses were performed, and graphs were constructed, using Prism version 11 (GraphPad Software, San Diego, CA, USA). The value of “n” refers to the number of cells for patch-clamp electrophysiology, Ca^2+^-imaging experiments, and STED experiments; the number of arteries for pressure myography; and the number of animals for ddPCR experiments. Statistical analyses were performed using unpaired Student’s *t* tests and one-way and two-way analysis of variance (ANOVA). A *P-*value < 0.05 was considered statistically significant for all analyses.

### Study approval

All animal care procedures and experimental protocols involving animals complied with the NIH Guide for the Care and Use of Laboratory Animals and were approved by the University Committee on Animal Resources (UCAR) at the University of Rochester and the Institutional Animal Care and Use Committee (IACUC) of the University of California, San Francisco.

## Data availability

The manuscript and the Supplemental Materials present all the data needed to evaluate the study’s conclusions. All original data sets and analyses, including individual comparisons, are also reported in the Supporting Data Values file.

## xxxx

**Supplemental Figure 1.** M**y**ogenic **tone is impaired in cerebral parenchymal arterioles from aged *Col4a1* mutant mice.** (**A**) Representative diameter recordings in isolated cerebral parenchymal arterioles from 12-month-old *Col4a1^+/+^* and *Col4a1^+/G394V^*mice during stepwise increases in intraluminal pressure from 5 to 140 mmHg. Active diameter traces are shown in black for *Col4a1^+/+^* controls and orange for *Col4a1^+/G394V^* arterioles. Passive diameter traces obtained in Ca^2+^-free PSS are shown in gray. Dashed lines indicate baseline diameter at 5 mmHg. (**B**) Summary data showing myogenic tone (%) as a function of intraluminal pressure. Data are presented as means ± SEM (n = 8 to 9 arterioles from 4 animals per group; \**P* < 0.05; 2-way ANOVA with Šídák’s multiple comparisons). (**C**) Representative diameter recordings in cerebral parenchymal arterioles from 12-month-old *Col4a1^+/+^*and *Col4a1^+/G1344D^* mice during stepwise increases in intraluminal pressure from 5 to 140 mmHg. Active diameter traces are shown in black for *Col4a1^+/+^* controls and red for *Col4a1^+/G1344D^* arterioles. Passive diameter traces obtained in Ca^2+^-free PSS are shown in gray. Dashed lines indicate baseline diameter at 5 mmHg. (**D**) Summary data showing myogenic tone (%) as a function of intraluminal pressure. Data are presented as means ± SEM (n = 7 to 17 arterioles from 4 to 8 animals per group; \**P* < 0.05; 2-way ANOVA with Šídák’s multiple comparisons).

**Supplemental Figure 2.** M**o**deling **the effects of segmental and combined loss of myogenic tone on downstream pressure transmission and capillary flow.** A simplified single-path resistance model was used to simulate the effects of segmental myogenic tone loss over an arterial pressure range of 70 to 140 mmHg. (**A**) Modeled capillary inlet pressure plotted as a function of arterial pressure under conditions of combined myogenic tone loss in pial arteries and parenchymal arterioles (PAs), loss of myogenic tone in PAs, loss of myogenic tone in pial arteries, and intact myogenic tone. (**B**) Relative capillary flow plotted over the same arterial pressure range and under the same conditions shown in panel A. Flow values are normalized to the intact-tone condition at 70 mmHg.

**Supplemental Figure 3.** S**T**OC **frequency and amplitude do not differ between SMCs from 3-month-old *Col4a2^+/+^*and *Col4a2^+/G646D^* mice.** (**A**) Representative STOCs recorded from SMCs voltage-clamped at membrane potentials ranging from -60 to -10 mV. (**B**) Summary of STOC frequency as a function of membrane potential. (**C**) Summary of STOC amplitude as a function of membrane potential. Data are presented as means ± SEM (n = 6 or 7 cells from 4 animals per group; ns, not significant; 2-way ANOVA with Šídák’s multiple comparisons).

## Reference

1. Markus HS, and Joutel A. The pathogenesis of cerebral small vessel disease and vascular cognitive impairment. Physiol Rev. 2025;105(3):1075–171.

2. Pantoni L. Cerebral small vessel disease: from pathogenesis and clinical characteristics to therapeutic challenges. Lancet Neurol. 2010;9(7):689–701.

3. Joutel A, and Faraci FM. Cerebral small vessel disease: insights and opportunities from mouse models of collagen IV-related small vessel disease and cerebral autosomal dominant arteriopathy with subcortical infarcts and leukoencephalopathy. Stroke. 2014;45(4):1215–21.

4. Joutel A, Haddad I, Ratelade J, and Nelson MT. Perturbations of the cerebrovascular matrisome: A convergent mechanism in small vessel disease of the brain? J Cereb Blood Flow Metab. 2016;36(1):143–57.

5. Gould DB, Phalan FC, Breedveld GJ, van Mil SE, Smith RS, Schimenti JC, et al. Mutations in Col4a1 cause perinatal cerebral hemorrhage and porencephaly. Science. 2005;308(5725):1167–71.

6. Gould DB, Phalan FC, van Mil SE, Sundberg JP, Vahedi K, Massin P, et al. Role of COL4A1 in small-vessel disease and hemorrhagic stroke. N Engl J Med. 2006;354(14):1489–96.

7. Lanfranconi S, and Markus HS. COL4A1 mutations as a monogenic cause of cerebral small vessel disease: a systematic review. Stroke. 2010;41(8):e513–8.

8. Boyce D, McGee S, Shank L, Pathak S, and Gould D. Epilepsy and related challenges in children with COL4A1 and COL4A2 mutations: A Gould syndrome patient registry. Epilepsy Behav. 2021;125:108365.

9. Massoudi D, Miner JH, and Gould DB. Collagen IV in Gould syndrome and Alport syndrome. Nat Rev Nephrol. 2025;21(11):778–93.

10. Bonaventura E, Marelli D, Carugo U, Politano D, Spaccini L, Parazzini C, et al. The Expanding Phenotypic Spectrum of COL4A1- and COL4A2-Related Disorders: A Systematic Review. Pediatr Neurol. 2026;183:10–21.

11. Gasparini S, Balestrini S, Saccaro LF, Bacci G, Panichella G, Montomoli M, et al. Multiorgan manifestations of COL4A1 and COL4A2 variants and proposal for a clinical management protocol. Am J Med Genet C Semin Med Genet. 2024;196(4):e32099.

12. Jeanne M, and Gould DB. Genotype-phenotype correlations in pathology caused by collagen type IV alpha 1 and 2 mutations. Matrix Biol. 2017;57-58:29–44.

13. Tambala D, Vassar R, Snow J, Balestrini S, Bersano A, Guey S, et al. COL4A1 and COL4A2-related disorders: Clinical features, diagnostic guidelines, and management. Genet Med. 2025;27(9):101514.

14. Whittaker E, Thrippleton S, Chong LYW, Collins VG, Ferguson AC, Henshall DE, et al. Systematic Review of Cerebral Phenotypes Associated With Monogenic Cerebral Small-Vessel Disease. J Am Heart Assoc. 2022;11(12):e025629.

15. Rannikmäe K, Davies G, Thomson PA, Bevan S, Devan WJ, Falcone GJ, et al. Common variation in COL4A1/COL4A2 is associated with sporadic cerebral small vessel disease. Neurology. 2015;84(9):918–26.

16. Traylor M, Persyn E, Tomppo L, Klasson S, Abedi V, Bakker MK, et al. Genetic basis of lacunar stroke: a pooled analysis of individual patient data and genome-wide association studies. Lancet Neurol. 2021;20(5):351–61.

17. Yamasaki E, Ali S, Sanchez Solano A, Thakore P, Smith M, Wang X, et al. Faulty TRPM4 channels underlie age-dependent cerebral vascular dysfunction in Gould syndrome. Proc Natl Acad Sci U S A. 2023;120(5):e2217327120.

18. Yamasaki E, Thakore P, Ali S, Sanchez Solano A, Wang X, Gao X, et al. Impaired intracellular Ca(2+) signaling contributes to age-related cerebral small vessel disease in Col4a1 mutant mice. Sci Signal. 2023;16(811):eadi3966.

19. Thakore P, Yamasaki E, Ali S, Sanchez Solano A, Labelle-Dumais C, Gao X, et al. PI3K block restores age-dependent neurovascular coupling defects associated with cerebral small vessel disease. Proc Natl Acad Sci U S A. 2023;120(35):e2306479120.

20. Davis MJ, Earley S, Li YS, and Chien S. Vascular mechanotransduction. Physiol Rev. 2023;103(2):1247–421.

21. Earley S, Waldron BJ, and Brayden JE. Critical role for transient receptor potential channel TRPM4 in myogenic constriction of cerebral arteries. Circ Res. 2004;95(9):922–9.

22. Gonzales AL, Yang Y, Sullivan MN, Sanders L, Dabertrand F, Hill-Eubanks DC, et al. A PLCγ1-dependent, force-sensitive signaling network in the myogenic constriction of cerebral arteries. Sci Signal. 2014;7(327):ra49.

23. Reading SA, and Brayden JE. Central role of TRPM4 channels in cerebral blood flow regulation. Stroke. 2007;38(8):2322–8.

24. Jeanne M, Labelle-Dumais C, Jorgensen J, Kauffman WB, Mancini GM, Favor J, et al. COL4A2 mutations impair COL4A1 and COL4A2 secretion and cause hemorrhagic stroke. Am J Hum Genet. 2012;90(1):91–101.

25. Jeanne M, Jorgensen J, and Gould DB. Molecular and Genetic Analyses of Collagen Type IV Mutant Mouse Models of Spontaneous Intracerebral Hemorrhage Identify Mechanisms for Stroke Prevention. Circulation. 2015;131(18):1555–65.

26. Dabertrand F, Krøigaard C, Bonev AD, Cognat E, Dalsgaard T, Domenga-Denier V, et al. Potassium channelopathy-like defect underlies early-stage cerebrovascular dysfunction in a genetic model of small vessel disease. Proc Natl Acad Sci U S A. 2015;112(7):E796–805.

27. Wenceslau CF, McCarthy CG, Earley S, England SK, Filosa JA, Goulopoulou S, et al. Guidelines for the measurement of vascular function and structure in isolated arteries and veins. Am J Physiol Heart Circ Physiol. 2021;321(1):H77–H111.

28. Borgstrom A, Hauert B, Kappel S, Zoni E, Kiener M, Stoklosa P, et al. Small Molecular Inhibitors Block TRPM4 Currents in Prostate Cancer Cells, with Limited Impact on Cancer Hallmark Functions. J Mol Biol. 2021;433(17):166665.

29. Gonzales AL, Amberg GC, and Earley S. Ca2+ release from the sarcoplasmic reticulum is required for sustained TRPM4 activity in cerebral artery smooth muscle cells. Am J Physiol Cell Physiol. 2010;299(2):C279–88.

30. Nelson MT, Cheng H, Rubart M, Santana LF, Bonev AD, Knot HJ, et al. Relaxation of arterial smooth muscle by calcium sparks. Science. 1995;270(5236):633–7.

31. Knot HJ, Standen NB, and Nelson MT. Ryanodine receptors regulate arterial diameter and wall [Ca2+] in cerebral arteries of rat via Ca2+-dependent K+ channels. J Physiol. 1998;508 ( Pt 1):211–21.

32. Perez GJ, Bonev AD, Patlak JB, and Nelson MT. Functional coupling of ryanodine receptors to KCa channels in smooth muscle cells from rat cerebral arteries. J Gen Physiol. 1999;113(2):229–38.

33. Cheranov SY, and Jaggar JH. Sarcoplasmic reticulum calcium load regulates rat arterial smooth muscle calcium sparks and transient K(Ca) currents. J Physiol. 2002;544(Pt 1):71–84.

34. Lukyanenko V, Viatchenko-Karpinski S, Smirnov A, Wiesner TF, and Györke S. Dynamic regulation of sarcoplasmic reticulum Ca(2+) content and release by luminal Ca(2+)-sensitive leak in rat ventricular myocytes. Biophys J. 2001;81(2):785–98.

35. Baddeley D, Jayasinghe I, Lam L, Rossberger S, Cannell MB, and Soeller C. Optical single-channel resolution imaging of the ryanodine receptor distribution in rat cardiac myocytes. Proc Natl Acad Sci U S A. 2009;106(52):22275–80.

36. Pritchard HAT, Pires PW, Yamasaki E, Thakore P, and Earley S. Nanoscale remodeling of ryanodine receptor cluster size underlies cerebral microvascular dysfunction in Duchenne muscular dystrophy. Proc Natl Acad Sci U S A. 2018;115(41):E9745–e52.

37. Xie Y, Yang Y, Galice S, Bers DM, and Sato D. Size Matters: Ryanodine Receptor Cluster Size Heterogeneity Potentiates Calcium Waves. Biophys J. 2019;116(3):530–9.

38. Mejía-Alvarez R, Kettlun C, Ríos E, Stern M, and Fill M. Unitary Ca2+ current through cardiac ryanodine receptor channels under quasi-physiological ionic conditions. J Gen Physiol. 1999;113(2):177–86.

39. Rajagopal V, Bass G, Walker CG, Crossman DJ, Petzer A, Hickey A, et al. Examination of the Effects of Heterogeneous Organization of RyR Clusters, Myofibrils and Mitochondria on Ca2+ Release Patterns in Cardiomyocytes. PLoS Comput Biol. 2015;11(9):e1004417.

40. Krishnan V, Ali S, Gonzales AL, Thakore P, Griffin CS, Yamasaki E, et al. STIM1-dependent peripheral coupling governs the contractility of vascular smooth muscle cells. Elife. 2022;11.

41. Hell SW, and Wichmann J. Breaking the diffraction resolution limit by stimulated emission: stimulated-emission-depletion fluorescence microscopy. Opt Lett. 1994;19(11):780–2.

42. Hernández Mesa M, van den Brink J, Louch WE, McCabe KJ, and Rangamani P. Nanoscale organization of ryanodine receptor distribution and phosphorylation pattern determines the dynamics of calcium sparks. PLoS Comput Biol. 2022;18(6):e1010126.

43. Porter VA, Bonev AD, Knot HJ, Heppner TJ, Stevenson AS, Kleppisch T, et al. Frequency modulation of Ca2+ sparks is involved in regulation of arterial diameter by cyclic nucleotides. Am J Physiol. 1998;274(5):C1346–55.

44. Wellman GC, Santana LF, Bonev AD, and Nelson MT. Role of phospholamban in the modulation of arterial Ca(2+) sparks and Ca(2+)-activated K(+) channels by cAMP. Am J Physiol Cell Physiol. 2001;281(3):C1029–37.

45. Ullrich ND, Valdivia HH, and Niggli E. PKA phosphorylation of cardiac ryanodine receptor modulates SR luminal Ca2+ sensitivity. J Mol Cell Cardiol. 2012;53(1):33–42.

46. Wehrens XH, Lehnart SE, Reiken S, Vest JA, Wronska A, and Marks AR. Ryanodine receptor/calcium release channel PKA phosphorylation: a critical mediator of heart failure progression. Proc Natl Acad Sci U S A. 2006;103(3):511–8.

47. Glass DB, Cheng HC, Mende-Mueller L, Reed J, and Walsh DA. Primary structural determinants essential for potent inhibition of cAMP-dependent protein kinase by inhibitory peptides corresponding to the active portion of the heat-stable inhibitor protein. J Biol Chem. 1989;264(15):8802–10.

48. Li Y, Baylie RL, Tavares MJ, and Brayden JE. TRPM4 channels couple purinergic receptor mechanoactivation and myogenic tone development in cerebral parenchymal arterioles. J Cereb Blood Flow Metab. 2014;34(10):1706–14.

49. Hartmann DA, Coelho-Santos V, and Shih AY. Pericyte Control of Blood Flow Across Microvascular Zones in the Central Nervous System. Annu Rev Physiol. 2022;84:331–54.

50. Ratelade J, Klug NR, Lombardi D, Angelim M, Dabertrand F, Domenga-Denier V, et al. Reducing Hypermuscularization of the Transitional Segment Between Arterioles and Capillaries Protects Against Spontaneous Intracerebral Hemorrhage. Circulation. 2020;141(25):2078–94.

51. Dixon RE, Navedo MF, Binder MD, and Santana LF. Mechanisms and physiological implications of cooperative gating of clustered ion channels. Physiol Rev. 2022;102(3):1159–210.

52. Galice S, Xie Y, Yang Y, Sato D, and Bers DM. Size Matters: Ryanodine Receptor Cluster Size Affects Arrhythmogenic Sarcoplasmic Reticulum Calcium Release. J Am Heart Assoc. 2018;7(13).

53. Shen X, van den Brink J, Bergan-Dahl A, Kolstad TR, Norden ES, Hou Y, et al. Prolonged β-adrenergic stimulation disperses ryanodine receptor clusters in cardiomyocytes and has implications for heart failure. Elife. 2022;11.

54. Bovo E, Huke S, Blatter LA, and Zima AV. The effect of PKA-mediated phosphorylation of ryanodine receptor on SR Ca(2+) leak in ventricular myocytes. J Mol Cell Cardiol. 2017;104:9–16.

55. Asghari P, Scriven DR, Sanatani S, Gandhi SK, Campbell AI, and Moore ED. Nonuniform and variable arrangements of ryanodine receptors within mammalian ventricular couplons. Circ Res. 2014;115(2):252–62.

56. Vizurraga A, Adhikari R, Yeung J, Yu M, and Tall GG. Mechanisms of adhesion G protein-coupled receptor activation. J Biol Chem. 2020;295(41):14065–83.

57. Leon K, Cunningham RL, Riback JA, Feldman E, Li J, Sosnick TR, et al. Structural basis for adhesion G protein-coupled receptor Gpr126 function. Nat Commun. 2020;11(1):194.

58. Paavola KJ, Sidik H, Zuchero JB, Eckart M, and Talbot WS. Type IV collagen is an activating ligand for the adhesion G protein-coupled receptor GPR126. Sci Signal. 2014;7(338):ra76.

59. Kakogiannos N, Scalise AA, Martini E, Maderna C, Benvenuto AF, D’Antonio M, et al. GPR126 is a specifier of blood-brain barrier formation in the mouse central nervous system. J Clin Invest. 2024;134(15).

60. Favor J, Gloeckner CJ, Janik D, Klempt M, Neuhäuser-Klaus A, Pretsch W, et al. Type IV procollagen missense mutations associated with defects of the eye, vascular stability, the brain, kidney function and embryonic or postnatal viability in the mouse, Mus musculus: an extension of the Col4a1 allelic series and the identification of the first two Col4a2 mutant alleles. Genetics. 2007;175(2):725–36.

61. Kuo DS, Labelle-Dumais C, Mao M, Jeanne M, Kauffman WB, Allen J, et al. Allelic heterogeneity contributes to variability in ocular dysgenesis, myopathy and brain malformations caused by Col4a1 and Col4a2 mutations. Hum Mol Genet. 2014;23(7):1709–22.

