## Supplemental Figures for "Ryanodine receptor cluster remodeling links *COL4A2* mutation to cerebral artery dysfunction"

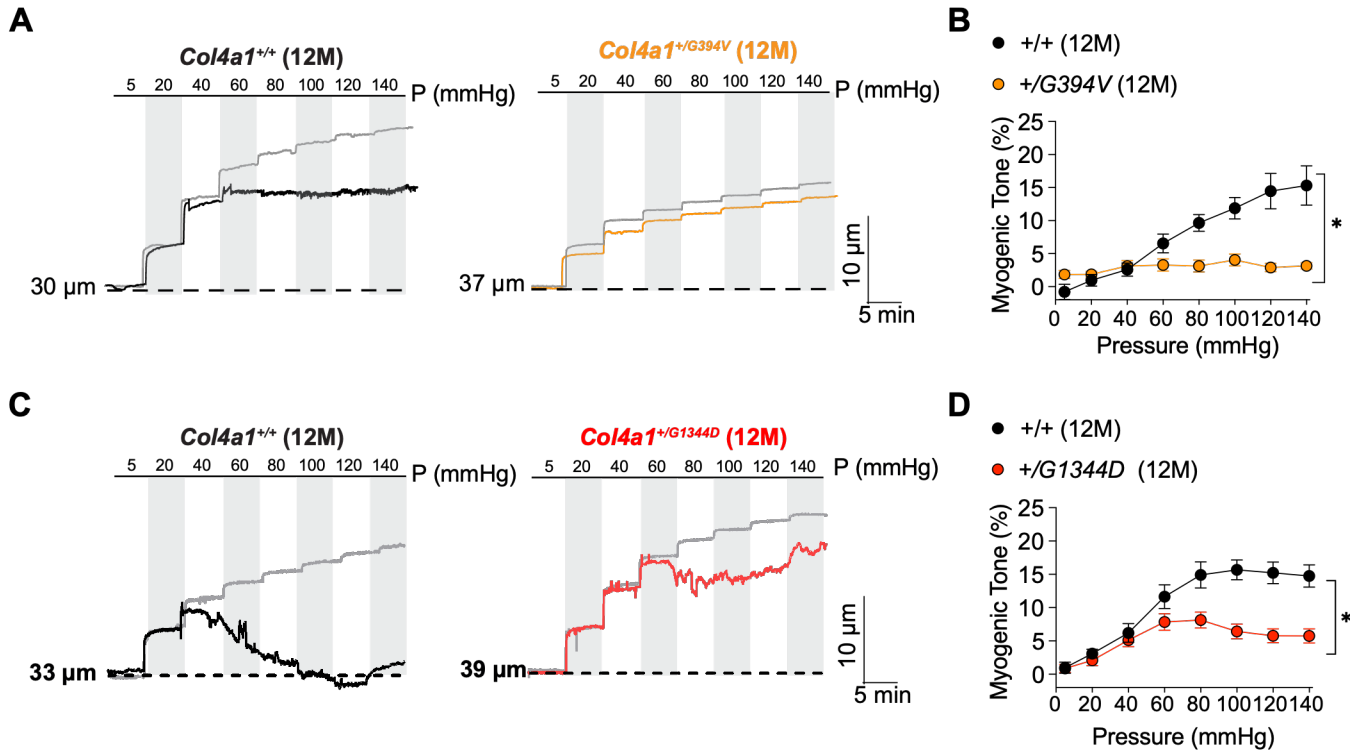

**Supplemental Figure 1. Myogenic tone is impaired in cerebral parenchymal arterioles from aged *Col4a1* mutant mice.** (A) Representative diameter recordings in isolated cerebral parenchymal arterioles from 12-month-old *Col4a1*<sup>+/+</sup> and *Col4a1*<sup>+/G394V</sup> mice during stepwise increases in intraluminal pressure from 5 to 140 mmHg. Active diameter traces are shown in black for *Col4a1*<sup>+/+</sup> controls and orange for *Col4a1*<sup>+/G394V</sup> arterioles. Passive diameter traces obtained in Ca<sup>2+</sup>-free PSS are shown in gray. Dashed lines indicate baseline diameter at 5 mmHg. (B) Summary data showing myogenic tone (%) as a function of intraluminal pressure. Data are presented as means ± SEM (n = 8 to 9 arterioles from 4 animals per group; \**P* < 0.05; 2-way ANOVA with Šidák's multiple comparisons). (C) Representative diameter recordings in cerebral parenchymal arterioles from 12-month-old *Col4a1*<sup>+/+</sup> and *Col4a1*<sup>+/G1344D</sup> mice during stepwise increases in intraluminal pressure from 5 to 140 mmHg. Active diameter traces are shown in black for *Col4a1*<sup>+/+</sup> controls and red for *Col4a1*<sup>+/G1344D</sup> arterioles. Passive diameter traces obtained in Ca<sup>2+</sup>-free PSS are shown in gray. Dashed lines indicate baseline diameter at 5 mmHg. (D) Summary data showing myogenic tone (%) as a function of intraluminal pressure. Data are presented as means ± SEM (n = 7 to 17 arterioles from 4 to 8 animals per group; \**P* < 0.05; 2-way ANOVA with Šidák's multiple comparisons).

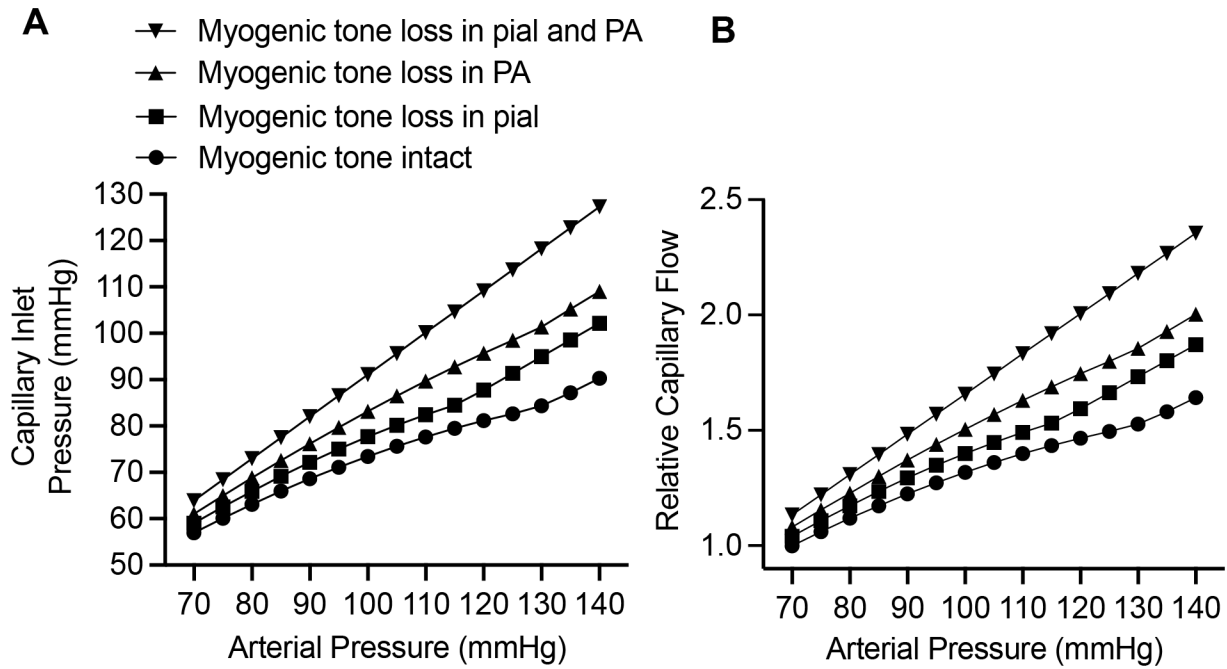

**Supplemental Figure 2. Modeling the effects of segmental and combined loss of myogenic tone on downstream pressure transmission and capillary flow.** A simplified single-path resistance model was used to simulate the effects of segmental myogenic tone loss over an arterial pressure range of 70 to 140 mmHg. **(A)** Modeled capillary inlet pressure plotted as a function of arterial pressure under conditions of combined myogenic tone loss in pial arteries and parenchymal arterioles (PAs), loss of myogenic tone in PAs, loss of myogenic tone in pial arteries, and intact myogenic tone. **(B)** Relative capillary flow plotted over the same arterial pressure range and under the same conditions shown in panel A. Flow values are normalized to the intact-tone condition at 70 mmHg.

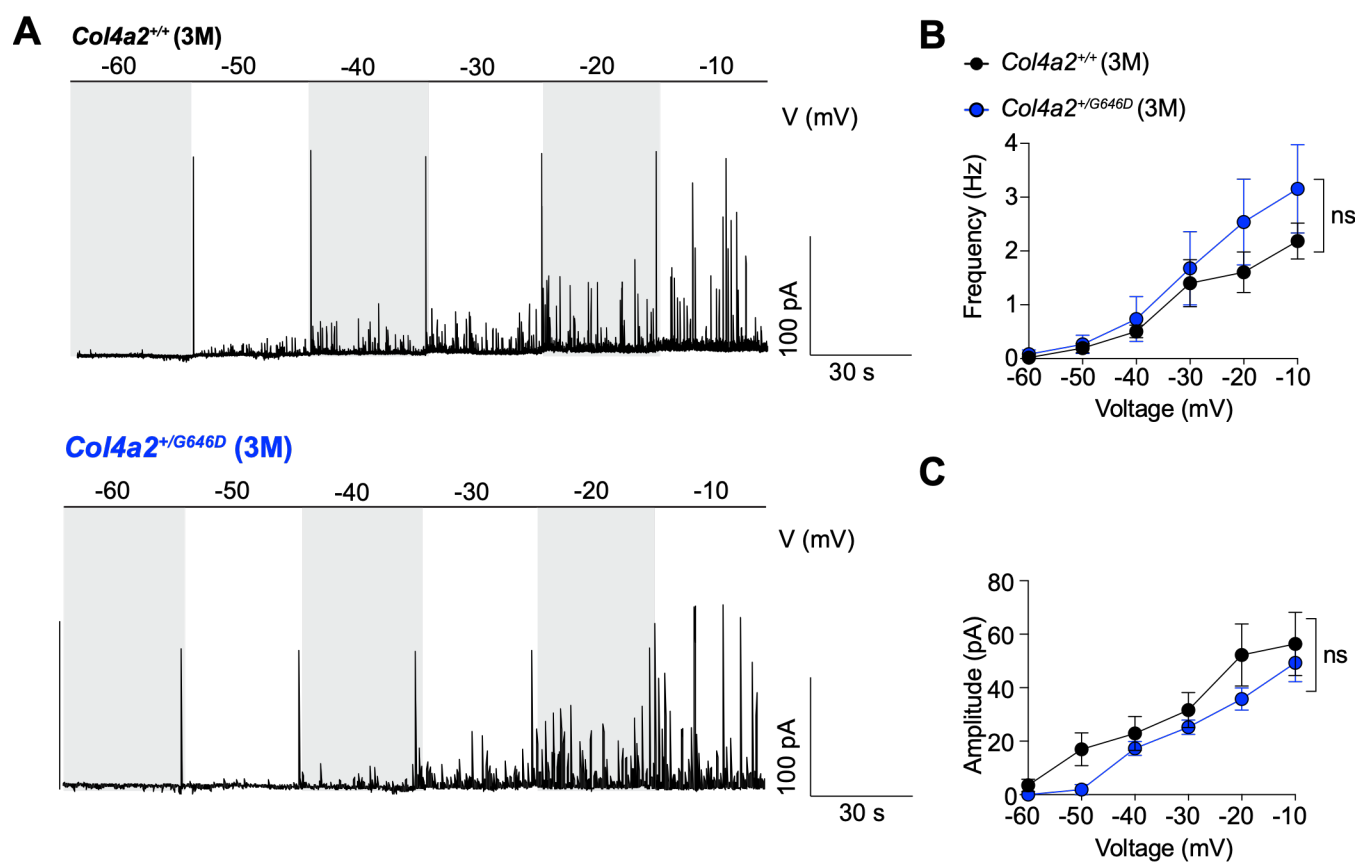

**Supplemental Figure 3. STOC frequency and amplitude do not differ between SMCs from 3-month-old *Col4a2*<sup>+/+</sup> and *Col4a2*<sup>+/G646D</sup> mice.** (A) Representative STOCs recorded from SMCs voltage-clamped at membrane potentials ranging from -60 to -10 mV. (B) Summary of STOC frequency as a function of membrane potential. (C) Summary of STOC amplitude as a function of membrane potential. Data are presented as means ± SEM ($n = 6$ or $7$ cells from $4$ animals per group; ns, not significant; 2-way ANOVA with Šídák's multiple comparisons).
